# Ligase-mediated programmable genomic integration (L-PGI): an efficient site-specific gene editing system that overcomes the limitations of reverse transcriptase-based editing systems

**DOI:** 10.1101/2024.09.27.615478

**Authors:** Angela Xinyi Nan, Michael Chickering, Christopher Bartolome, Neeta Shadija, Dan Li, Brett Estes, Jessica Von Stetina, Wei Li, Jason Andresen, Jesse C Cochrane, Chen Bai, Jason Gatlin, Jie Wang, Davood Norouzi, Sandeep Kumar, Maike Thamsen Dunyak, Leonard Chavez, Anmol Seth, Shakked Halperin, Jonathan D Finn, Jenny Xie

## Abstract

Since their discovery, CRISPR/Cas9 systems have been repurposed for programmable targeted genomic editing. This has led to unprecedented advancement of gene editing for therapeutic benefit. Initial uses of CRISPR/Cas9 were focused on gene disruption via DNA cleavage, but significant engineering led to systems for single base editing as well as insertion, deletion and manipulation of short stretches of genomic sequences using nicking Cas9 and RT-based methods. These technologies allowed safer and more precise editing but were limited to small corrections and showed significantly reduced efficiencies in nondividing cells, presenting difficulty for translation to *in vivo* therapies. To find an alternate editing strategy that could address these shortcomings, we revisited the mechanism of DNA nicking by nCas9. nCas9 nicking creates a free 5’ phosphate group and a 3’ hydroxyl group on the complementary strand of the target sequence. Under ordinary conditions in the cell these ends are re-joined by endogenously expressed ligases to repair DNA back to wild-type. If, however, a DNA fragment containing the desired edit were present, ligation of the nicked genomic DNA with the delivered fragment could result in gene editing. We demonstrate that optimization of each component and introduction of a chemically modified high affinity splinting DNA allows a variety of ligase-based edits, including longer edits not efficient with RT-based systems, at high efficiencies and fidelities that minimize genomic byproducts in both dividing and nondividing cells as well as *in vivo* in adult mice. Here we present the first therapeutically relevant ligation-based programmable gene editing technology, L-PGI.

---

CRISPR/Cas9 revolutionized the controlled manipulation of genetic information in eukaryotic organisms due to its ease of retargeting, requiring only modifying a portion of the RNA sequence of CRISPR/Cas9 to match the genomic location intended to be edited [1, 2]. Cas9 was suited to gene disruption as DNA cleavage resulted in double stranded breaks (DSBs). However, deleterious side effects including heterogenous on target edit outcomes containing insertions or deletions (indels), potential off target cleavage, chromosomal rearrangements and even chromosomal loss, discouraged its use for some genetic diseases [3]. Engineering of the CRISPR/Cas9 system led to tools with greater finesse, such as single strand nicking (nCas9) or complete catalytic inactivation (dCas9) [2]. dCas9 coupled with deaminases led to enzymes that can make specific single nucleotide changes [4, 5]. nCas9 combined with writing enzymes, such as a Moloney murine lentiviral reverse transcriptase (MMLV-RT, RT) led to enzymes that could write new sequence to introduce small corrections instead of simply disrupting the target site [6].

When nCas9 nicks the genome, it leaves behind a 3’ hydroxyl group [6]. This moiety can be extended by an RT but is also a suitable substrate for DNA ligase, which under ordinary conditions simply re-joins the 5’ and 3’ free ends and reverts the nick back to the endogenous sequence. If a synthetic DNA template containing alternate sequence were present, this action could result in gene insertion. Pre-synthesizing the corrective edit as a complete strand of DNA would simplify the corrective gene editing to a single step and remove the uncertainty of multi-step writing fidelity and endogenous dNTP availability of RT-based systems [7]. We address this by designing the DNA donor as a single stranded DNA and introducing a nonintegrating splinting DNA (splint) component to create a new method of gene editing: ligase-mediated programmable genomic integration (L-PGI).

L-PGI addresses many of the problems observed in RT-based editing methods. Namely, the low efficiencies of long edits (>10 base pairs (bp)) as well as the poor fidelity and efficiency seen in non-dividing cells, likely due to the decreased intracellular dNTP pools in cells not actively replicating DNA [7]. The L-PGI system consists of nCas9, DNA ligase, a ligase-mediating single guide RNA with a 3’ extension (lmgRNA), a single stranded DNA (ssDNA) donor, and a ssDNA splint. The splint contains three hybridization domains that bind to the donor, the guide RNA, and the genomic target site. These three interactions serve two key functions: efficient delivery of the donor via nCas9 ribonucleoprotein (RNP) nuclear localization and alignment of the donor to the nicked genomic flap for ligation. Similar to prime editing (PE), the inclusion of a nicking guide (ngRNA) targeting the opposite strand approximately 50 – 100 bp away from the edit site improves efficiency, and is employed for all experiments comparing the two editing methods. We also opted to develop L-PGI using only mRNA and synthetic guide formats and focus on nondividing cells to minimize translational risk.

Through chemical and sequence optimization of each oligonucleotide component to improve affinity, stability, and activity, we report efficiencies of small corrections (1 – 3 nucleotide changes) in therapeutically relevant loci of up to 50% in HEK293T cells and 70% in primary human hepatocytes (PHH) with up to 99% fidelity (edits lacking any indels). Pairing two complexes together (paired L-PGI, pL-PGI), we observe up to 82% total efficiency and 63% fidelity for placement of a 38 bp Bxb1 integrase recognition site (attB) in PHH. Using pL-PGI to place attB leads to 15% on target integration of 30 kb of transgene cargo when co-delivering integrase and helper-dependent adenovirus template (HDAd), representing 60% usage of available attBs. pL-PGI demonstrates good translatability across important model species *in vitro*, resulting in similar or significantly superior efficiency and fidelity compared to PE. Lastly, we present mouse data showing 7.9% attB placement using pL-PGI compared to 2.5% by dual PE, demonstrating that the advantages of L-PGI seen *in vitro* carry over *in vivo*. In summary, due to its good tolerability, high potency, and favorable fidelity as well as compatibility with therapeutically relevant delivery modalities such as lipid nanoparticles (LNP), we believe L-PGI has the potential to expand the gene editing medicine toolbox and enable treatment of previous inaccessible genetically driven diseases.

## Results

### L-PGI Mechanism and Optimization

Ligase-mediated programmable genomic integration (L-PGI) capitalizes on nicking of the non-target strand of the genome by nCas9 leaving a free hydroxyl group on the 3’ DNA flap [6]. This free 3’ DNA end can now be a substrate for DNA ligase and the genome can be directly modified using a donor DNA sequence containing a free 5’ phosphate. After ligation, incorporation of the free 3’ end of the donor into the genome is aided by addition of a short homology arm (10 – 20 nt) downstream of the edit. Using a complementary oligonucleotide splint that contains a donor binding site (DBS), flap (or genomic) binding site (FBS) and a guide binding site (GBS) to link the donor, genome and 3’ end of a single guide RNA, all the editing components can be localized to enable efficient gene editing (Fig. 1a). In its final configuration, L-PGI consists of nCas9 and ligase-mediating guide RNA (lmgRNA) to nick the genome, splint DNA to facilitate the insertion of donor DNA and a DNA ligase to ligate the junctions. The newly ligated donor DNA competes with the endogenous strand and successful incorporation results in the desired editing outcome (Fig. 1b) [6]. Using a Cy5-labeled dsDNA substrate in an *in vitro* biochemical assay, we showed that inclusion of all the L-PGI components was necessary for formation of a product corresponding to ligation of the donor DNA into the substrate (Lane 3, Fig. 1c).

**Figure 1.**
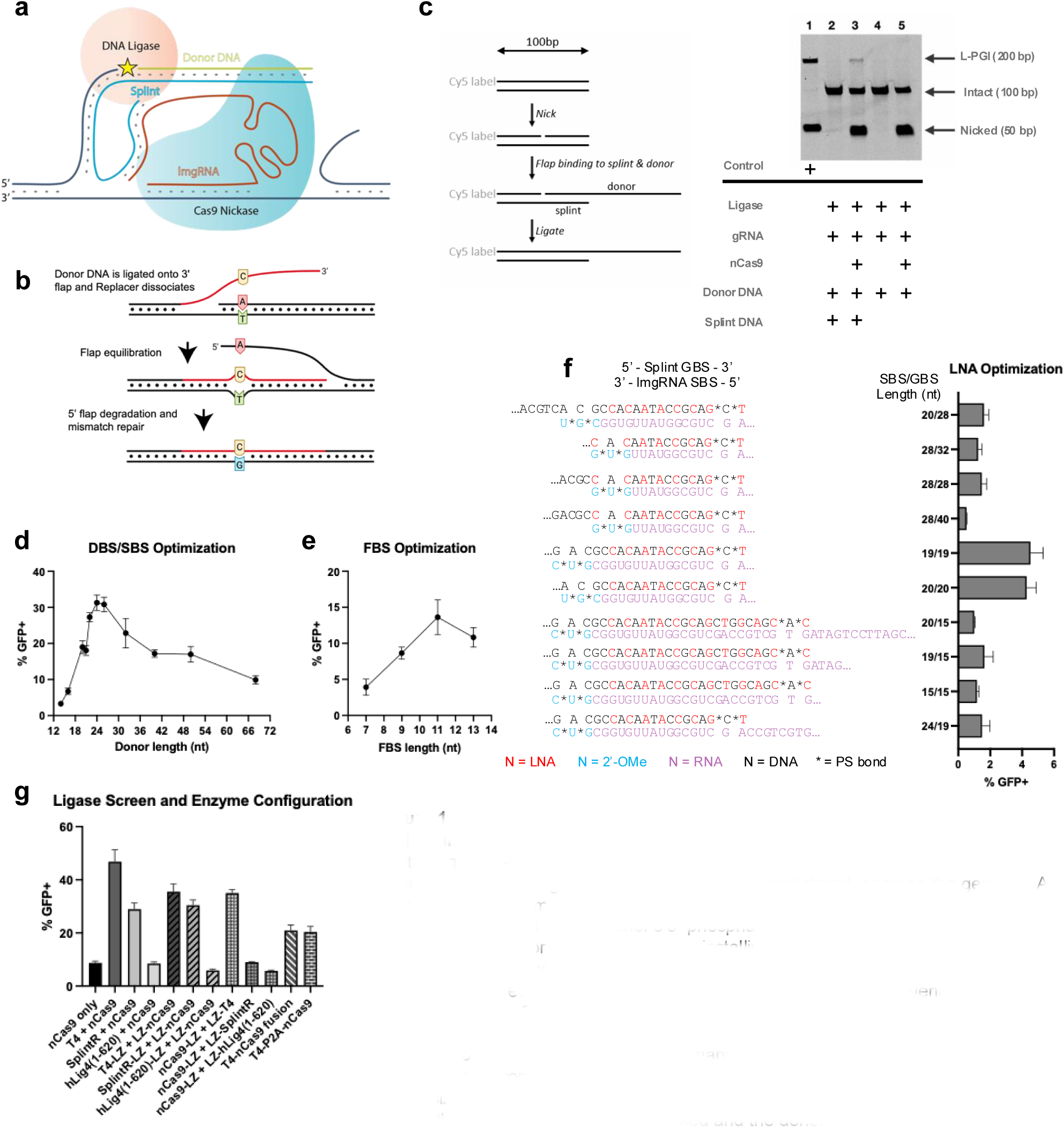
Overview of L-PGI editing mechanism, biochemical proof of concept, and optimizations in HEK293T GFP reporter cell line. (A) Schematic diagram illustrating the L-PGI editing complex. H840A Cas9 nickase (nCas9) is guided to a protospacer by the lmgRNA and nicks the non-target strand, opening the genome. A splint bound to the lmgRNA and donor DNA then hybridizes to the exposed 3’ flap of the genome, positioning the donor’s 5’ phosphate at the OH of the 3’ flap. Ligase is able to join the donor DNA to the 3’ flap, installing the edit in the genome. (B) Example of incorporation of the L-PGI edit for an A to C transversion after ligation. The donor DNA (shown in red) contains the desired edit with homologous sequence arms to the left and the right of the edit. After ligation, the donor competes with the endogenous strand for genome incorporation via the mismatch repair (MMR) pathway. (C) Validation of edit strategy using recombinant protein in vitro. A 100 bp 5’-Cy5 labeled DNA substrate containing the protospacer and PAM is mixed with chemically synthesized lmgRNA, splint, and donor DNA, as well as nCas9 and T4 ligase proteins. During the reaction, the subtrate is nicked and the donor DNA is ligated to the remaining 5’-Cy5 fragment while they are both bound to the splint. The donor DNA contains a PAM mutation to prevent repeat nicking by nCas9. After 1 hour at 37C, reaction products are denatured and visualized with PAGE. Lane 1 contains 50 nt and 200 nt DNA fragments as positive controls representing the expected edit products. Lanes 2 to 5 are L-PGI reactions containing components as labeled below. The 200-nt ligation product is present only when all components are included (lane 3). (D) Optimization of DBS/SBS length using the HEK293T GFP reporter cell line. (E) Optimization of FBS length using the HEK293T reporter cell line. (F) Effect of splint GBS and lmgRNA SBS lengths on BFP to GFP conversion efficiency. Nucleotide sequences are shown as they are bound to each other and in some cases a part of the GBS or SBS functions as a single stranded linker. All splints include the same DBS and FBS (not shown) and lmgRNAs include identical scaffolds and spacers (not shown). (G) Efficiencies of linked and split mRNA architecture testing various placements of leucine zippers (LZ), P2A, or XTEN linkers on nCas9 and either T4, SplintR, or human ligase IV (hLig4) truncations.

We then derived a HEK293T reporter cell line containing a blue fluorescent protein (BFP) that converts to green fluorescent protein (GFP) upon successful editing of 3 nucleotides at positions +2, +4, and +5 from the nick (Extended Data Fig. 1a) [8]. Initial experiments using an unmodified DNA splint did not yield detectable conversion from BFP to GFP. However, substituting locked nucleic acids (LNAs) in the GBS and DBS regions of the splint, thereby increasing the stability of the splint and its affinity for the donor and lmgRNA, led to low but detectable, levels of GFP (Extended Data Fig. 1b) [9]. We then systematically varied the lengths of the DBS, FBS and GBS and found that 24 – 28 nt, 11 – 13 nt and 19 nt resulted in the highest GFP for the three sequence components respectively (Fig. 1d-f). Additional optimization of the LNA position and number led to a splint design with LNAs alternating throughout the DBS and in the 3’ end of the GBS (Extended Data Fig. 1c). Addition of LNAs in the FBS did not improve efficiency (Extended Data Fig. 1d). Moving forward, we used a DNA splint containing a 19 nt GBS, 13 nt FBS, and 24 nt DBS to assess editing efficiency of L-PGI but locus-dependent optimization is expected to further improve the system for specific targets.

We next explored other avenues for improving the L-PGI system, including enzyme selection, mRNA architecture and IVT condition optimization. Initial work was conducted using T4 ligase as it is a well-characterized highly efficient DNA ligase [10]. When compared with other ligases, human DNA ligase IV (hLig4) and PBCV-1 DNA ligase (SplintR), T4 ligase consistently led to the highest GFP signal (Fig. 1g) [11]. We also investigated colocalization and fusion of nCas9 and ligase as these strategies had been successfully for other genome editing technologies [6]. However, linking nCas9 and T4 ligase through either a flexible peptide linker, self-cleaving peptide or leucine zippers (LZ) had either no effect or a detrimental effect on editing efficiencies (Fig. 1g) [12]. When LZ-based linking was used for SplintR, we found that LZ placement strongly impacted efficiency in an orientation-dependent manner (Extended Data Fig. 1e). Because of this we included LZ on the C terminus of ligase and N terminus of nCas9 for all designs. Finally, we found that including N1-methylpseudouridine during mRNAs production led to an almost 2-fold increase in GFP fluorescence compared to uridine, as has been previously reported (Extended Data Fig. 1f) [13]. Based on these findings, we used split LZ co-localized nCas9 and T4 ligase translated from N1-methylpseudouridine mRNAs for subsequent L-PGI target efficiency analysis.

Gene editing by L-PGI is dependent on all the components of the system. No editing was observed without nCas9, lmgRNA, donor, or splint, suggesting that nicking and both splint and donor are required. However, some editing was achieved when T4 ligase or phosphorylation of the 5’ end of donor DNA were omitted, suggesting that endogenous ligases involved in DNA repair can aid in L-PGI (Extended Data Fig. 1g). We demonstrate that the observed editing was not due to the splint acting as a template for HDR by comparing L-PGI oligonucleotides with an HDR donor designed for editing of BFP to GFP. Cas9 co-delivered with the HDR template produced ∼10% GFP conversion while no significant editing was observed with the splint and donor (Extended Data Fig. 1h).

### L-PGI For Therapeutic Targets

L-PGI can correct both point mutations in disease associated loci as well as introduce longer edits. We explored a set of disease relevant mutations in HEK293T and after extensive optimization of chemical modifications [14–16], enzyme architecture, and transfection conditions, were able to reach precise editing efficiencies of between 25% and 50% for three different loci with <1.5% indel generation for each edit (Extended data Fig. 2a-d). To enhance translatability into *in vivo* models and therapeutic applications, we focused subsequent work in non-dividing primary human hepatocytes (PHH) on liver associated genomic targets. To aid in the evaluation of the technology for therapeutic use, we also move to a metric for describing the quality of the edit, fidelity, defined as the proportion of sequencing reads aligned to the edit outcome that have the correct edit, with no insertions or deletions. L-PGI oligonucleotide components were designed based on high efficiency nCas9 spacers to install disease-relevant mutations in wild type cells including H1069Q in *ATP7B* for Wilson’s Disease and C282Y in *HFE* for Hemochromatosis [15, 17]. In PHH, L-PGI demonstrated a 10-fold advantage in editing efficiency over prime editing at *ATP7B*, reaching over 20% efficiency with 99% fidelity (Fig. 2a). The 2nt correction at *HFE* was performed with over 30% efficiency and similar fidelity (Fig. 2b).

**Figure 2.**
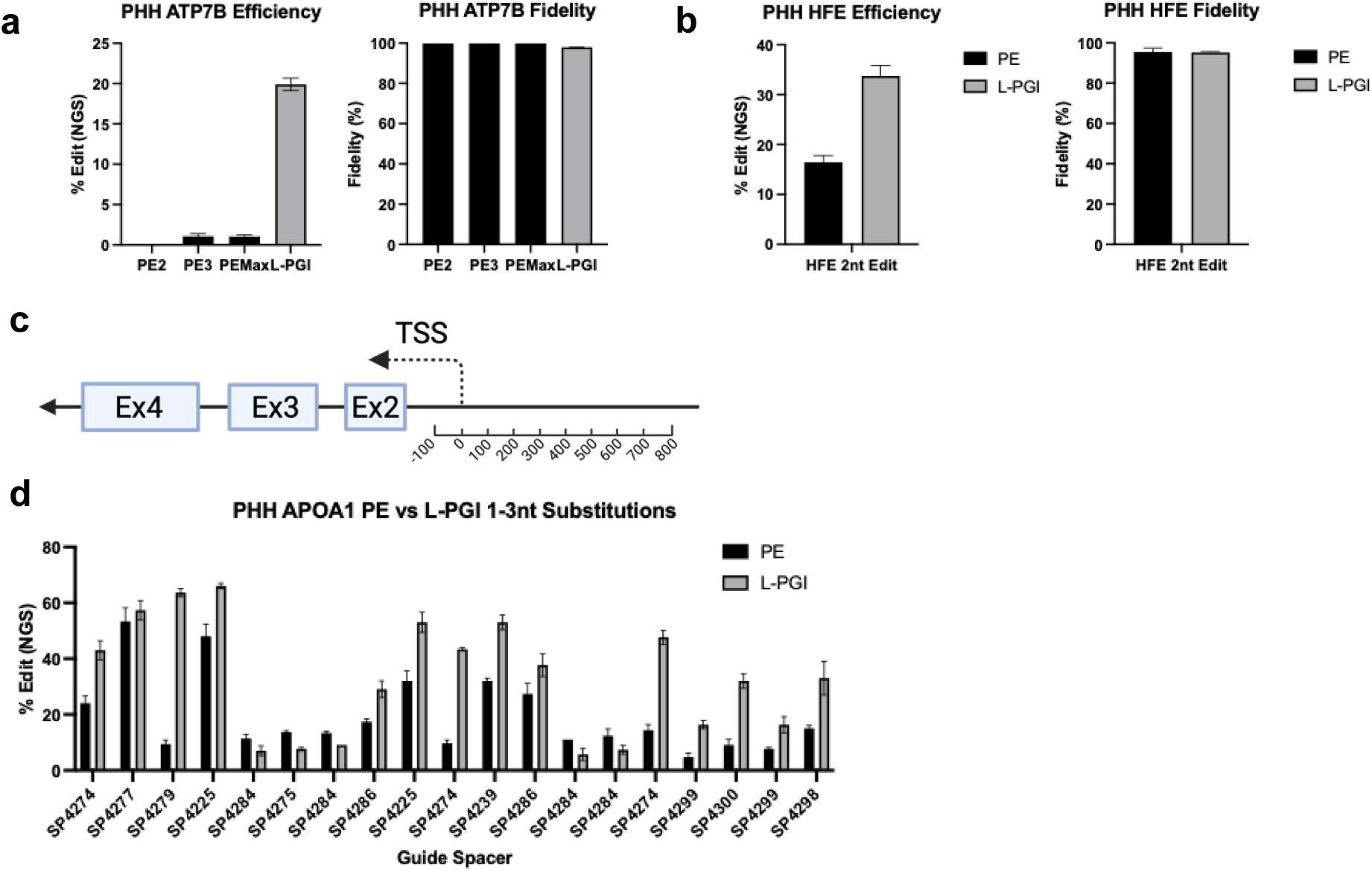
Comparison of prime editing (PE) versus L-PGI for installation of point mutations in 4 disease-relevant loci in PHH. (A) Efficiency and fidelity of 2 nt substitution of +5 G to T and +8 G to T to install H1069Q with silent mutation in *ATP7B* demonstrating therapeutic potential for correction of mutation to treat Wilson’s Disease. L-PGI was compared to PE2, PE3, and PEMax. (B) Efficiency and fidelity of 2 nt substitution of UGC to UAU to install C282Y in *HFE* to show potential treatment for Hemochromatosis. L-PGI is compared to PE which in this and all following cases refers to nCas9 fused to engineered RT. (C) Schematic map showing location of guide target window in a 900 bp range spanning the transcription start site (TSS) preceding exon 2 of *APOA1*. (D) Efficiency of 1 – 3 nt substitutions performed by either PE or L-PGI shown side by side using the same spacer sequences for either pegRNA or lmgRNA and ngRNA by Sanger Sequencing and ICE analysis.

We then explored the application of precision gene editing for modifying gene expression as an alternate therapeutic method for permanent upregulation of specific genes to treat disease. We selected *APOA1* for proof of concept of this strategy [18]. We selected spacers to perform 1 – 3 nt edits in a 900 bp window in the promotor region upstream of the transcription start site that would result in placement transcription factor binding motif-like sites to enhance expression (Fig. 2c). We found up to 70% efficiency with L-PGI compared to 35% with PE (Fig. 2d). Interestingly, though identical target spacers and nicking guides were used for both L-PGI and PE, L-PGI generally matched or outperformed PE. The pattern suggests that RT is more constrained by sequence identity while L-PGI can consistently approach the theoretical edit ceiling based on Cas9 targeting efficiency.

One of the advantages of L-PGI is that pre-synthesized DNA is used as the edit itself. By shifting the placement of the homology arm in the donor, we designed 14 bp motif edits either as a complete insertion or as a sequence substitution by simultaneously excising endogenous sequence of the same length (Fig. 3a). Using L-PGI, we found up to 35% total editing compared to ∼ 5% using RT-based (pegRNA) editing and superior fidelity with L-PGI (Fig. 3b). Insertion generally outperformed sequence substitution both by ligation and RT based methods likely due to the favorability of strand displacement closer to the nick site. Comparing the indel generation between L-PGI and PE, we find that L-PGI maintains a lower error rate across all small insertion types tested (Fig. 3c). The additional errors with PE appear in the edit itself as a combination of substitutions and indels and are likely due to erroneous RT activity (Fig. 3d). While substitution errors appear in L-PGI as well, these are potentially the result of error during chemical synthesis and should diminish with higher purity donors. We then tested longer insertions, including a 38 bp Bxb1 integrase attachment site (attB), and found up to 12% correct editing with L-PGI compared to only 0.5% with RT (Fig. 3e). Though the edit outcomes for attB insertion via L-PGI were still favorable compared to RT, we saw a reduction in efficiency consistent with the drop in efficiency observed in earlier GFP reporter HEK293T line using longer donor lengths. Overall, we determined that genome editing by L-PGI is not as limited by insertion size and that editing outcomes are generally of higher quality in nondividing cells compared to PE.

**Figure 3.**
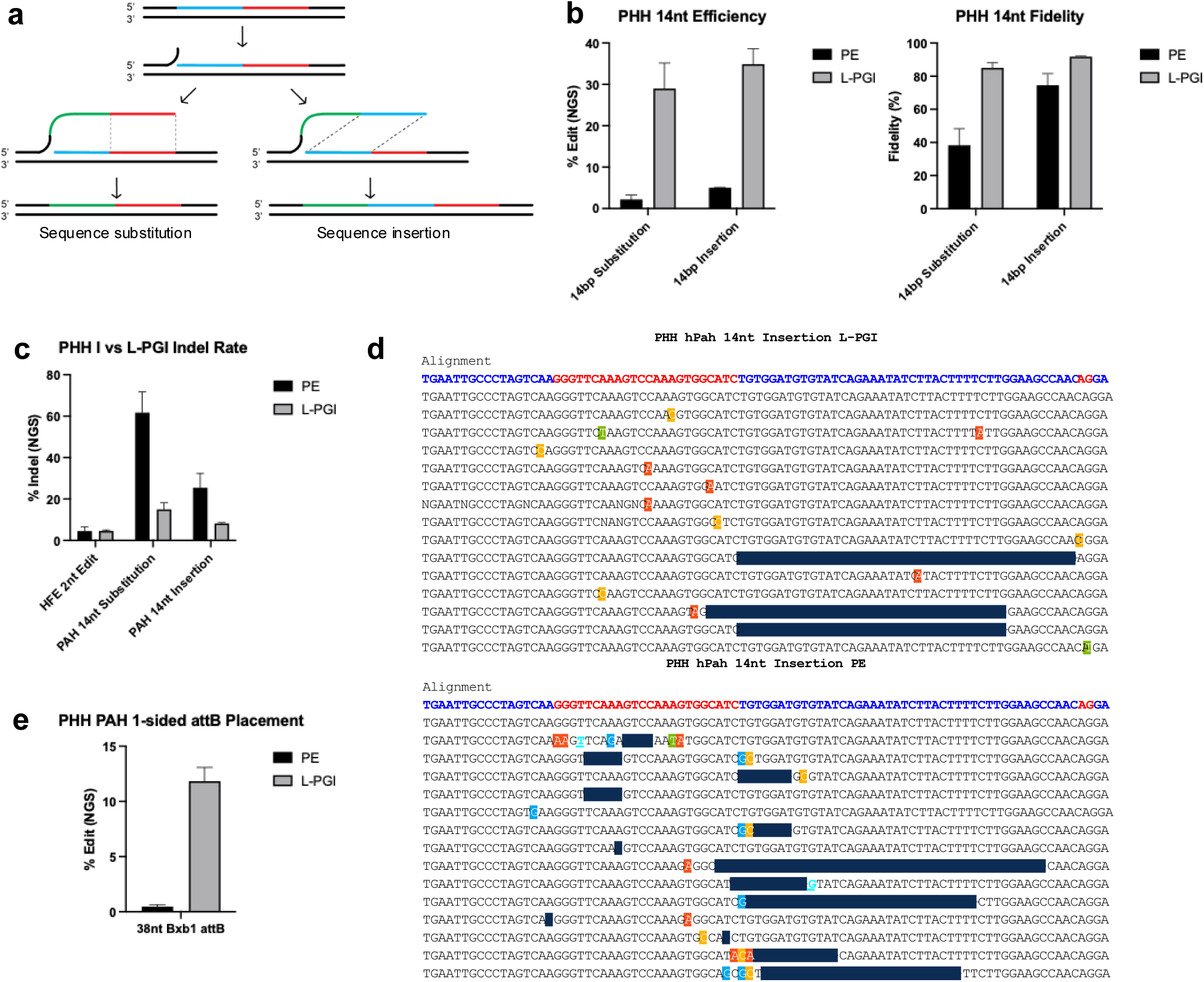
Comparison of prime editing (PE) versus L-PGI larger small corrections in PHH. (A) Illustration of the two methods of placing the 14 nt motif with and without excision of endogenous sequence representing end cases of editing resulting in either preserving endogenous sequence or reading frame. The endogenous sequence downstream of the nick is shown in blue and red. The donor contains the 14nt insertion followed by sequence homologous to the red for sequence replacement or to the blue for sequence insertion. (B) Highest total efficiencies and maximum fidelities observed using either sequence replacement or insertion observed using either PE or L-PGI in *PAH* intron 1 obtained through component ratio optimization. Efficiencies and fidelity vary depending on component ratio, for complete optimization data refer to Extended Data Figure 3. (C) Comparison of overall lowest indel generation rates between PE and L-PGI. Indel rate is calculated as the rate of erroneous deletions and insertions occurring within the nick to nick edit window without discrimination between size of error normalized to total edited reads. (D) Representative NGS alignment results showing top 15 reads each of 14 nt insertions in *PAH* intron 1 of PHH using L-PGI (top) and PE (bottom). Blacked out regions indicate deletions, highlighted bases show substitutions, and underlined bases contain insertions. In L-PGI the dominant errors are point substitutions in the donor region (24 nt in red in reference alignment) suggestive of error during chemical synthesis and excisions between the end of the donor and the nicking site on the opposite strand (2 nt in red) suggestive of error during heteroduplex resolution or flap excision during downstream DNA repair. In PE we find greater occurance of point substitutions and deletions in the edit window suggestive of RT transcription errors. (E) Comparison of 38 nt Bxb1 attB placement efficiency in *PAH* intron 1 using L-PGI or PE by NGS showing correct editing only.

### Paired L-PGI

Two L-PGI complexes (paired L-PGI, pL-PGI) simultaneously introducing edits on opposite strands of the genome could make sequence replacements or deletions with higher efficiencies than with a single complex (Fig. 4a). The lmgRNAs, donors and splints were designed so that the ligated flaps would be either complementary with variable hybridization length for sequence replacement, or complementary to the genome beyond the opposing nick for precise deletion (Fig. 4b). In both scenarios, the genomic region between the two nicks is excised by 5’ exonuclease degradation [19], resulting in a size change detectable by gel electrophoresis following target amplification (Extended Data Fig. 4a) [20, 21]. We used pL-PGI to delete the *C9ORF72* hexanucleotide repeat expansion in HEK293T cells (Extended Data Fig. 4b) and replace this 131 bp deleted region with a 38 bp Bxb1 attB site and confirmed this with gel electrophoresis (Extended Data Fig. 4c). We quantified the frequency of these edits using NGS amplicon sequencing and found that pL-PGI could install a 33 bp Pa01 attB site at *NOLC1* locus in HEK293T cells with ∼1.5 – 2 fold efficiency improvement over RT-based editing and ∼60% reduction in indels (Fig. 4c) [22]. In general the addition of second editing complex improves efficiency at the cost of increased indels though we find that pL-PGI results in the most favorable correct edit rate. pL-PGI was also used to delete 175 bp at the *VEGFA* locus in both HEK293T and PHH. In HEK293T, pL-PGI led to ∼50% deletion efficiency, compared to ∼20% for RT-based editors (Fig. 4d). In PHH, we saw ∼15% editing efficiency of the *VEGFA* locus with almost 100% fidelity (Fig. 4e).

**Figure 4.**
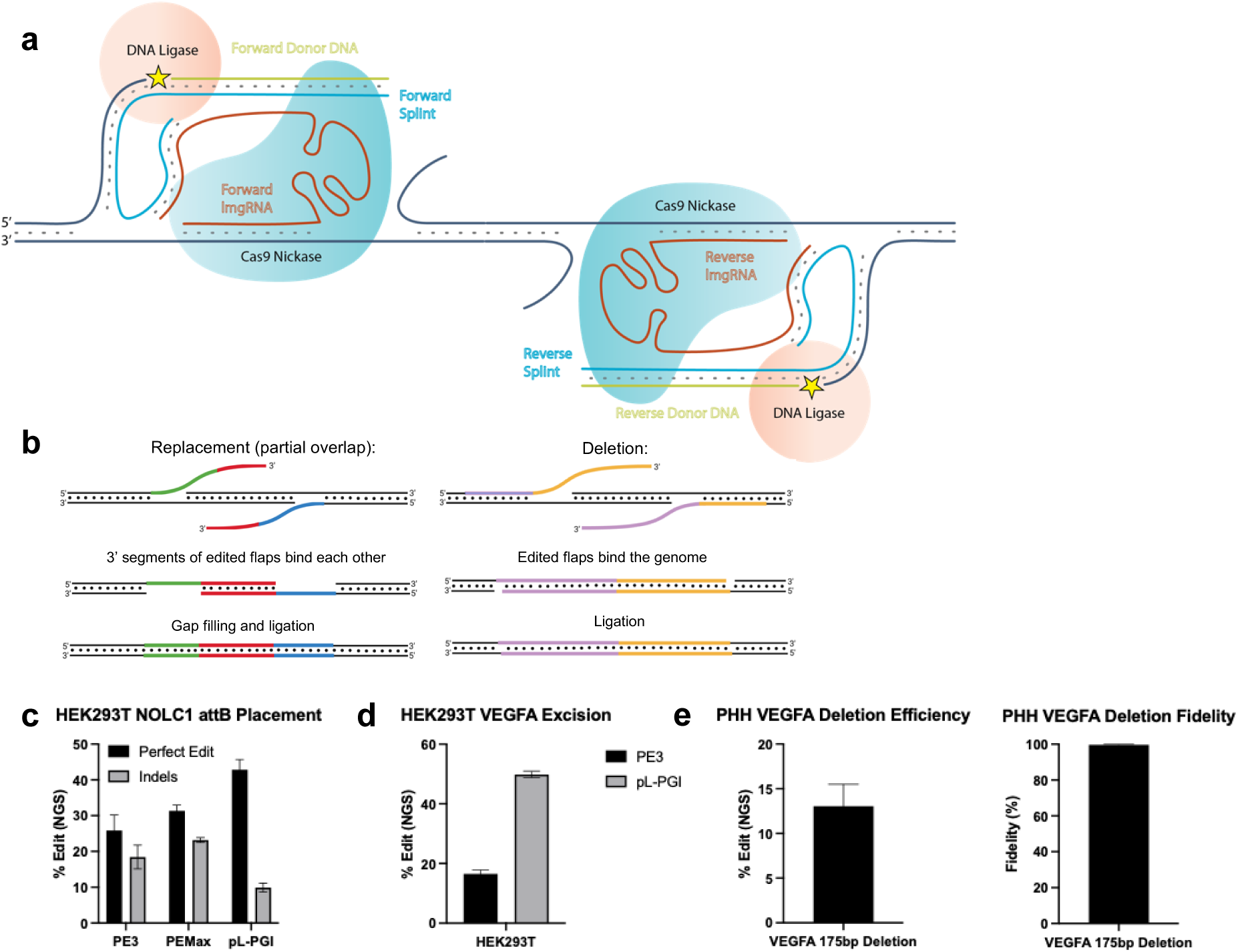
Paired L-PGI (pL-PGI) design and efficiencies in HEK293T for attB placement and excision. (A) Schematic diagram of two L-PGI complexes targeting opposite strands to install paired reverse complementary donors. (B) Design strategies demonstrating replacement type edit (left) by encoding the desired edit in reverse complementary partial overlap donors or deletion type edit (right) by having donors homologous to the genome flanking the sequence to be excised. (C) Pa01 33bp attB placement efficiencies using PE3, PEMax, or pL-PGI in *NOLC1* of HEK293T cells. (D) Efficiencies of 175 bp excision in *VEGFA* of HEK293T cells by either PE2 or pL-PGI. (E) Efficiency and fidelity of the same *VEGFA* deletion edit in PHH.

### Leveraging pL-PGI for Integrase Mediated Genomic Insertion

The data suggest that L-PGI and pL-PGI lead to more precise editing outcomes than prime editing likely because the entire edit is encoded in the donor DNA. This is a particularly attractive mechanism for carrying out longer inserts, including Bxb1 attB placement for integrase mediate gene insertion in non-dividing cells such as PHH [22–24]. Considering the additional chemically modified DNA components needed for pL-PGI, we first evaluated the cytotoxicity in PHH. Using an ATP-based cell viability assay, we found that pL-PGI reagents delivered with lipid-based transfection are well tolerated in PHH with minor decrease in viability at high doses of splints and donors (Extended Data Fig. 5a, b). pL-PGI relies on two reverse complementary donors that can encode a partial edit or a complete edit. We determined the optimal splint and donor architecture by symmetrically varying the length of overlap around the central dinucleotide while maintaining full complementarity between the splint and donor and found that the shortest overlap showed both best efficiency and fidelity (Fig. 5a). To minimize the impact of the attB secondary structure on editing efficiencies, we further explored the oligonucleotide lengths and overlaps and found we could improve efficiency for 20 bp overlap by decreasing the relative splint length but found that overall 10 bp overlap was still superior (Extended Data Fig. 5c). At the same time, we explored the ability of pL-PGI for placing Bxb1 attP attachment site instead of attB. We varied the donor overlaps for the 52 bp attP sequence in a similar manner and saw while fidelity was correlated with donor length, that efficiency was inversely related (Extended Data Fig. 5d). From our attB and attP insertion studies we concluded that the optimal editing efficiency and fidelity can be achieved with two donors with 10 bp overlap.

**Figure 5.**
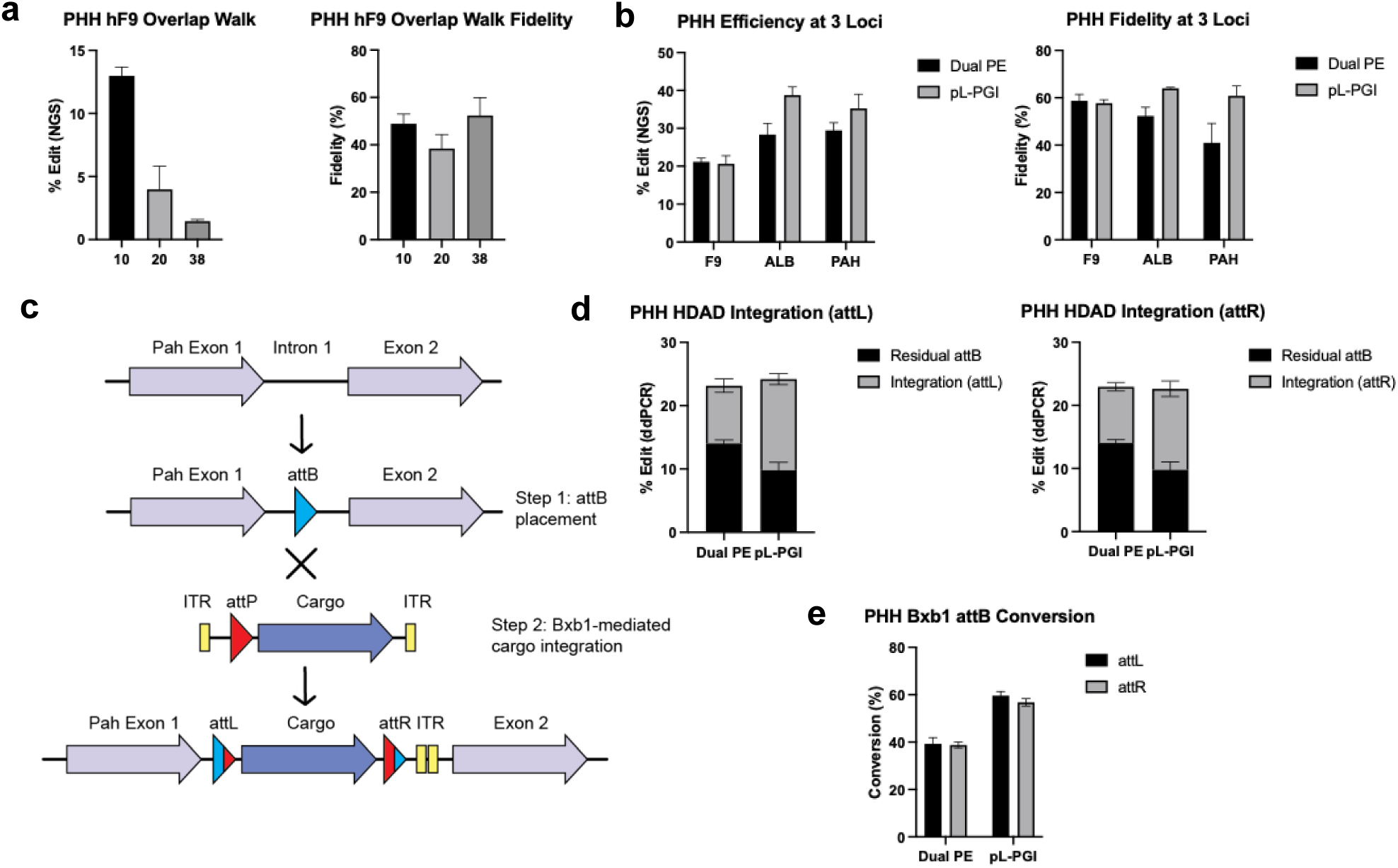
pL-PGI Bxb1 attB placement optimizations across therapeutic loci and implementation of pL-PGI for Bxb1-mediated integration in PHH. (A) Efficiency and fidelity of attB placement with symmetrically varying overlap for forward and reverse donors in F9. DBS and SBS were kept locked to maintain full donor splint hybridization and were 24, 29, 38 nt for 10,20, and 38 overlap respectively. (B) Application of 10 overlap pL-PGI for attB placement in *Factor 9 (F9)*, *Albumin (ALB)*, and *PAH* in PHH versus dual 20 overlap PE with fused nCas9 and engineered RT showing frequency of correct editing only. (C) Schematic illustration of 2-step attB placement followed by Bxb1-mediated integration of viral cargo showing expected genomic products. (D) Residual unconverted attB and total gene integration efficiency of helper-dependent adenovirus (HDAd) comparing attB placed by either dual PE or pL-PGI. Sum of both bars represents total edit efficiency for each attB placement method. (E) Comparison of attB conversion rate (calculated as ratio of integration to integration + residual attB) between dual PE and pL-PGI across both junctions. Comparable efficiencies and conversion rate across left and right junctions suggest successful integration of entrie intact 30 kb cargo.

Taking our lead overlap architecture into additional therapeutically relevant loci, we compared pL-PGI with 10 overlap against dual PE with 20 overlap at *F9*, *ALB*, and *PAH* [25–27]. We found 20 – 40% total editing efficiencies with pL-PGI, making it as efficient or more efficient than RT based editing at all loci tested (Fig. 5b). The fidelity of RT-based editing appears to be more locus dependent than pL-PGI whereby full 38 bp insertions are made with consistently with higher quality. Taken together, these findings suggest that pL-PGI is a viable method to install longer edits such as integrase landing site placement and perform deletions with higher efficiency and improved fidelity over prime editing.

The donor oligonucleotides are highly modified including 5’-methylation of all cytosines to enhance affinity and 3’ phosphorothioate bonds to protect them from nuclease degradation. However, these modifications may also interfere with gene integration by serine recombinases after installation into host genomes. In 2-step PGI, an attB is first placed in the host genome then co-delivery of Bxb1 and cargo carrying the corresponding attP result in integrase mediated gene insertion (Fig. 5c) [22–24]. Initial efforts using fully methylated cytosine donors for integrase-mediated gene insertion found low attB conversion. We then screened additional chemical modifications for attB placement in PHH and found that removal of all methylation while retaining two phosphorothioate bonds resulted in preservation of donor stability, edit efficiency and improved Bxb1 activity in the cell (Extended Data Fig. 5e, f). Using this modification pattern, we were able to install 38 bp attB into intron 1 of *PAH* and in the presence of Bxb1 mRNA and helper-dependent adenoviral (HDAd) DNA cargo (Extended Data Fig. 5g), measured over 15% whole gene insertion and 60% attB conversion by ddPCR (Fig. 5d, e, Extended Data Fig. 5h). We found higher attB conversion with pL-PGI than RT-based editing, consistent with the improvements in fidelity seen with pL-PGI.

### Adapting pL-PGI for *in vivo* gene editing

Having optimized L-PGI and pL-PGI in human cells, we took steps to move these systems *in vivo*. Using appropriately designed lmgRNAs, donors and splints we were able to efficiently install 38 bp attBs into intron 1 of the *PAH* gene in primary mouse hepatocytes (PMH) and primary cynomolgus hepatocytes (PCH), which are important surrogate cell types for non-human animal models. For the RT-based control in all experiments, we employed a fusion of nCas9 with engineered RT and 20 overlap dual attachment guide RNAs (atgRNAs) (dual PE) containing the same targeting sequences as pL-PGI. Comparing pL-PGI to optimized dual PE attB insertion, we observed similar correct and total edit efficiency in both PCH and PHH (Fig. 6a, b). However, in PMH we observed almost no correct insertion using dual PE while pL-PGI was able to achieve ∼11% placement of functional attBs (Fig. 6c). pL-PGI and dual PE have comparable fidelity in PCH, but pL-PGI has 1.5-fold higher fidelity in PHH, and a 10-100-fold gain in PMH (Fig. 6d-f). Additionally, pL-PGI and dual PE lead to different imperfect editing outcomes (Extended Data Fig. 6a, b). pL-PGI generally results in insertions containing the entire forward or reverse donor with blunt end joining to the second nick site. RT-based insertion shows partially written sequence or incorrect sequence that may be due to incorrect reverse transcription or to partial strand degradation among other possibilities. pL-PGI has a fidelity advantage over prime editing due to delivery of a pre-synthesized chemically protected DNA edit over *in situ* error-prone RT-based writing.

**Figure 6.**
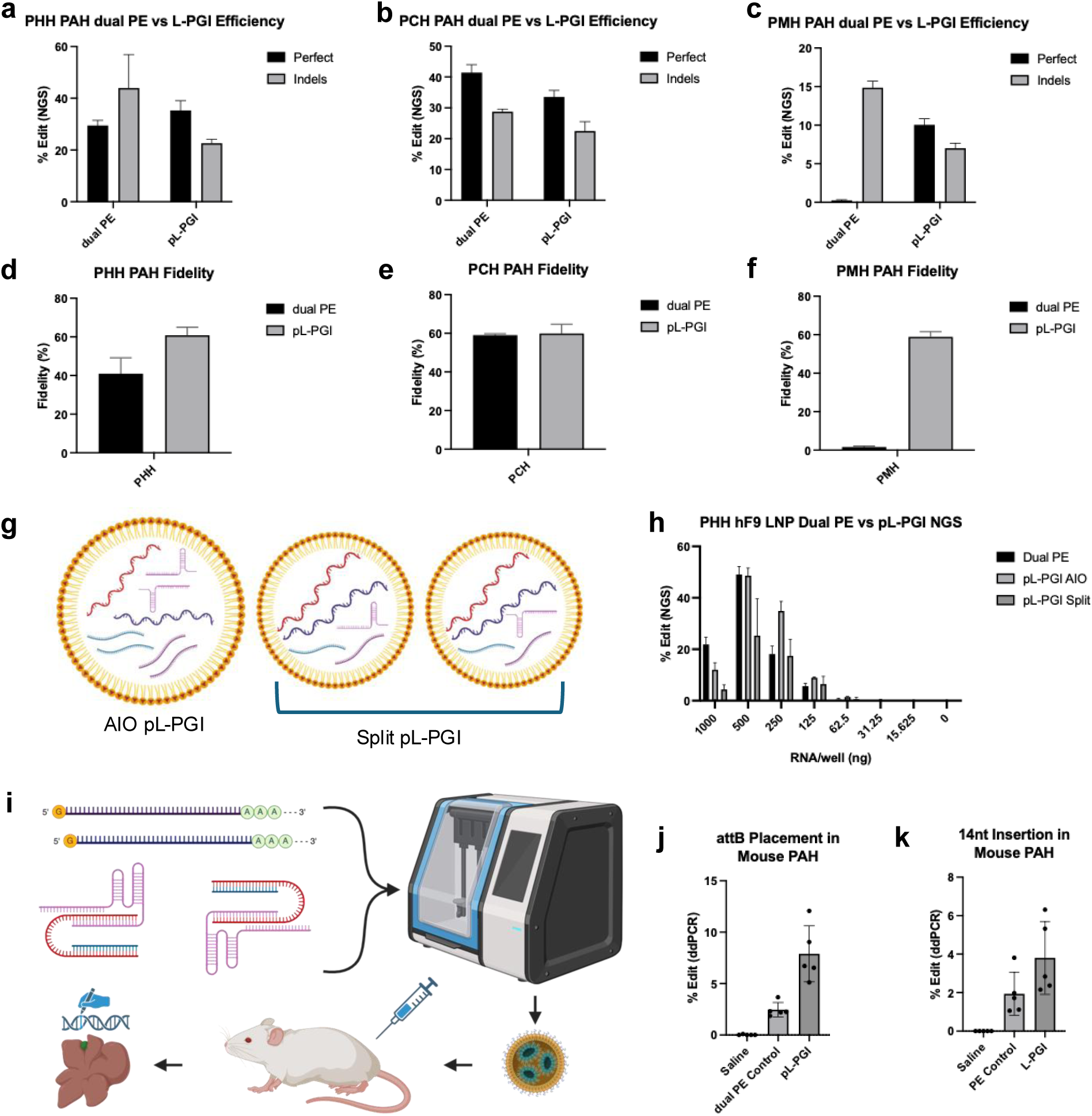
Translation of pL-PGI across primary cell species in vitro, delivery with lipid nanoparticles (LNP) in vitro, and efficiency in mice. (A-C) Perfect edit and indel efficiencies at PAH intron 1 in PHH, PCH, and PMH respectively using guides targeting the identical nicking sites for both edit methods in each comparison. (D-F) Fidelity comparison between pL-PGI and dual PE for the same edits in PHH, PCH, and PMH respectively, calculated by taking the percent of perfect beacons out of total edits. (G) pL-PGI LNP formulation payload schematic depicting all in one (AIO) delivery with one particle containing nCas9 mRNA, ligase mRNA, forward lmgRNA, forward splint, forward donor, reverse lmgRNA, reverse splint, and reverse donor or split delivery with two particles each containing both mRNAs and all forward or all reverse guide and oligonucleotide components. (H) Total edit potency of dual PE AIO, pL-PGI AIO, and pL-PGI split LNP targeting *F9* in PHH at different doses of LNP. (I) Schematic diagram of formulation and injection process in mouse, illustrated for pL-PGI with 8 components in AIO formulation. (J) Dual PE vs pL-PGI for BP in mouse *PAH* by ddPCR showing significant improvement in efficiency for pL-PGI vs dual PE demonstrating in vitro to in vivo translatability of PMH findings. (K) PE vs L-PGI for insertion of 14 nt edits in mouse *PAH* showing similar gain in efficiency with L-PGI.

Using pL-PGI for therapeutic indications also involves moving to higher purity reagents and formulation of the components in a relevant delivery modality such as lipid nanoparticles (LNP). We tested HPLC-purified synthetic human *F9* lmgRNAs and found a 1.5-fold improvement over crude lmgRNAs which translated to an overall ∼3-fold improvement in attB placement efficiency compared to RT-based attB placement with purified synthetic atgRNAs (Extended Data Fig. 6c). We also tested the effect of purification on donors and found improvement to 80% total editing with 64% fidelity at *PAH* in PHH (Extended Data Fig. 6d). Next, we compared single all in one (AIO) LNP and split LNP packaging designs for attB placement in *F9*, theorizing that splitting the formulation may minimize unwanted cross-hybridization between oligonucleotide components (Fig. 6g). We found that overall efficiencies improved but relative efficiencies using LNP delivery correlated well with MessengerMAX, adding confidence to the translatability of pL-PGI through delivery formats (Fig. 6h). Due to similar performance between AIO and split formulation methods, AIO was selected for *in vivo* formulation for simplicity. Lastly, we formulated mouse *F9* reagents into LNPs and found 10-fold higher attB placement in PMH with pL-PGI than dual PE, also consistent with earlier MessengerMAX results (Extended Data Fig. 6e).

Based on *in vitro* optimization, we selected the lead L-PGI 14 nt insertion and pL-PGI attB placement architecture and compared their efficiency against PE and dual PE respectively. High purity guide RNAs were synthesized in-house for pL-PGI and dual PE and other synthetic oligonucleotide components were provided by an external vendor (IDT) without purification. mRNAs and oligonucleotide components were co-formulated AIO and dosed to adult mice (Fig. 6i). Treatments were well tolerated and animals were sacrificed 1 week after dosing for analysis by ddPCR. We found up to 12.1% in one animal with an average of 7.9% attB placement by pL-PGI and an average of 2.5% by dual PE (Fig. 6j). For 14 nt insertion using all unpurified standard desalted oligonucleotides, we find L-PGI to result in an average of 3.8% insertion and PE 1.9% (Fig. 6k). Extrapolation from PMH *in vitro* would suggest that these efficiency differences are primary fidelity driven. We were encouraged by these results and anticipate that improvements of deliverable quality through synthesis improvement and formulation optimization would result in future efficiency gains.

In summary, we have demonstrated that L-PGI is functional in a range of dividing and non-dividing cells, can perform a range of edit classes (insertions, deletions, replacements), has increased potency and fidelity vs current RT-based writing systems (especially in non-dividing cells), and demonstrates good tolerability *in vivo* using clinically relevant LNP delivery.

## Discussion

Large precise insertions using CRISPR/Cas9 based systems have been difficult using the available gene editing systems [1, 28–30]. Here we describe L-PGI and pL-PGI, two new technologies that capitalize on the biochemistry of CRISPR/Cas9 nicking to ligate edits using fully synthetic DNA donors.

We first demonstrate that L-PGI and pL-PGI are capable of a broad range of edits in cycling cells and primary hepatocytes. Small edits (point mutations) can be installed to replace common founder mutations for diseases such as hereditary hemochromatosis (HH), with improved efficiency over RT-based systems especially for sequences difficult for RT to process. We have also demonstrated that L-PGI has significantly higher efficiency and fidelity vs RT systems for medium sized (∼14 bp) edits. This class of edit is essential for applications such as endogenous gene augmentation via installation of transcription factor binding sites. This advantage can also be applied to expand the edit range which reduces PAM availability restriction in Cas9 targeting. While we have not yet explored the maximum size limit of the L-PGI system, the ability to simultaneously delete and insert sequences potentially opens the door to exon replacements, which could treat a multitude of genetic diseases caused by repeat expansions [31].

We then demonstrate through a series of optimizations in primary mouse, human and cynomolgus hepatocytes, that L-PGI and pL-PGI are efficient methods for enabling large gene insertion via writing of integrase landing sites. pL-PGI leads to more accurate editing outcomes than RT-based integrase landing site insertion, and pL-PGI in combination with Bxb1 integrase inserts the *F9* coding gene (30 kb) in the *PAH* locus with significantly higher efficiency. Due to its capacity for longer insertions without compromising efficiency, pL-PGI can alternatively place 52 bp attP for Bxb1 integration. As integrases are not cargo size limited, this technology can enable precise and directed whole gene replacement that restores function under endogenous gene regulation [23].

Lastly, we explore the translational potential of L-PGI by evaluating the impact of reagent purity on potency *in vitro*, the feasibility of formulating 8 separate RNA and DNA components in a single LNP, and the efficiencies of L-PGI versus RT-based editing for 14 nt insertion and pL-PGI for 38 nt attB placement in adult mice. We find L-PGI to perform very favorably in both edit types *in vivo* even with unpurified reagents and expect optimizations in cargo quality and formulation to yield greater efficiencies in the future.

We consider the presented body of work to demonstrate that L-PGI is an effective and versatile gene editing system for *in vivo* liver therapeutics with advantages in the described key use cases. While we did not look at off targets in this work, we do not anticipate significant off target effects due to the inability of unbound donors to effectively access genomes and expect refinement of donor synthesis and purification processes to reduce error rates of on target edits. Future directions include expansion of the editing capabilities of L-PGI to perform efficiently for larger insertions through engineering of enzyme and oligonucleotide components and application of L-PGI for novel therapeutic strategies. Due to its success even in species challenging for RT-based editing such as mouse, L-PGI may have greater universal efficacy and could be coupled with delivery advancements for future therapies targeting important organs such as brain and spinal cord. By offering solutions to fundamental shortcomings of RT-based editing, we believe L-PGI will play an important role in the next generation of gene editing therapies.

## Methods and Materials

### *In vitro* transcription of mRNA

All mRNAs were generated via in vitro transcription (IVT) reactions using the HiScribe T7 High Yield RNA Synthesis Kit (New England Biolabs). Coding sequences were ordered as gBlocks (IDT) and cloned into an IVT vector that contains a single copy of the 5’UTR and two copies of the 3’UTR from the human beta globin gene, in addition to a 152nt polyA tail. Plasmid DNA containing coding sequences were linearized using an XbaI restriction site located immediately downstream of the polyA tail. Linearized plasmids were then purified via phenol:chloroform extraction followed by ethanol precipitation. mRNAs were produced via IVT reactions that contain Uridine-Triphosphate or N1-Methylpseudouridine-5’-Triphosphate (TriLink BioTech), and capped co-transcriptionally with CleanCap Reagent AG (3’ OMe) (TriLink BioTech). IVT reactions were incubated at 37°C for 2 hours, followed by DNAse I digestion of the template DNA. mRNA products were purified using LiCl precipitation, quantified (Qubit Fluorometric Quantification; ThermoFisher), and checked for integrity by denaturing gel electrophoresis.

### Preparation of synthetic oligonucleotides

Splints and donor DNAs were ordered from IDT as 100 nmole DNA oligos in IDTE buffer. The ssODN used for HDR was ordered from IDT as a 4nmole Ultramer DNA Oligo. lmgRNAs, pegRNAs, and atgRNAs were ordered from IDT as Custom Alt-R gRNAs, 10 nmol with standard desalting. lmgRNAs and atgRNAs were purified by HPLC. Nicking gRNAs were ordered from Synthego as synthetic sgRNA, 5 nmol. Synthego sgRNAs included Synthego’s standard scaffold and chemical modification pattern while IDT gRNAs were custom specified. Splints and donor DNAs were annealed together in NEBuffer 2 at 2μM by heating to 95C for 2 minutes and ramping down to 25C over a period of 30 minutes.

### *In vitro* biochemical assays

*In vitro* reactions were performed by first incubating lmgRNA (30nM final) and annealed splint and donor DNA (30nM final) with recombinant S. pyogenes nicking Cas9 (nCas9; IDT; 30nM final) for 10min at room temperature, followed by the addition of T4 ligase (NEB; 200U final), ATP (1mM final), and 5’-Cy5-labeled dsDNA substrate (3nM Final). Reactions were carried out in the presence of NEB Buffer 3.1 (1x final) at 37C for 1hr (final volume of 10ul). Reactions were terminated by the addition of 0.5% SDS and 100ug/ml Proteinase K, and incubated at 37C for 30min. Reaction products were then combined with 2x formamide gel loading buffer (90% formamide; 10% glycerol; 0.01% bromophenol blue), denatured at 95 °C for 10min, and separated by denaturing urea PAGE gel (15% TBE-urea, 55 °C, 200 V). DNA products were visualized by Cy5 fluorescence signal using a LI-COR Odyssey CLx imager.

### General cell culture conditions

HEK293T cells were purchased from ATCC and cultured in DMEM (11965092, Gibco) with 10% FBS (A3160501, Gibco). Cells were dissociated with 0.25% Trypsin-EDTA (15400054, Gibco) and seeded in 96-well PDL-coated tissue culture plates (354210, Corning) at a density of 25k cells per well. Cryopreserved primary human hepatocytes (HMCPMS Hu8403, Hu8449, Hu8450, Gibco) were recovered in Cryopreserved Hepatocyte Recovery Media (CM7000, Gibco) and plated at 42k cells per well in Hepatocyte Plating Media (A1217601, CM300, Gibco) on Collagen I coated 96 well plates (354407, Corning). Cryopreserved primary cynomolgus monkey hepatocytes (MKCP10 CY427, Gibco) were recovered in Hepatocyte Plating Media and plated to 48k cells per well. Cryopreserved primary mouse hepatocytes (MSCP10 MC945, Gibco) were recovered in Hepatocyte Plating Media and plated to 20k cells per well. 8 hours post recovery primary hepatocytes were washed and cultured in maintenance media with fresh media given again at 24 hours (CM400, A1217601, A2737501, Gibco).

### Generation of BFP+ HEK293T cell line

A lentiviral vector encoding BFP was transfected into HEK293T cells with VSV-G and psPAX2 helper plasmids. Supernatant was collected after 48h, centrifuged, and run through a 0.45-μm filter. The filtered supernatant was used to infect HEK293T cells in serial dilutions, and a well that contained ∼20% GFP+ cells was sorted by flow cytometry (BD FACSAria III, BD Biosciences) to result in a stable BFP+ HEK293T cell line with the majority of cells containing a single proviral integrant.

### Adenovirus-associated virus (AAV) production

Cargo was designed as self-complementary AAV containing Bxb1 attP attachment site, splice acceptor, and codon optimized human PAH exon 2-13 coding sequence with HiBit reporter tag (Promega) and cloned into plasmid backbone with Gibson assembly. AAV was produced by triple transfection with LK03 rep/cap and helper plasmid (Aldevron SF058826) in 293AAV Cell line (Cell Biolabs AAV-100). Virus was treated with PEG and purified with Iodixanol gradient centrifugation. Yield was determined by ITR assay titer using ddPCR.

### Helper-dependent adenovirus (HDAd) production

Cargo was designed containing Bxb1 attP attachment site, splice acceptor, and codon optimized human F9 exon 2-8 coding sequence and assembled into in Ad5 gutless plasmid backbone. Cargo was linearized and transfected in HEK293 116 Cre+ cells (Baylor University) followed by helper virus transduction. HDAd was propagated by serial coinfection and purified by CsCl ultracentrifugation. Yield was determined by GFP reporter assay titer using ddPCR.

### HEK293T transfection

After 16-24h, the medium was changed to OptiMEM (31985070, Thermo Fisher Scientific) and the cells were transfected at approximately 70% confluency. A transfection mix for a single well included 0.4 uL of Lipofectamine 2000 (11668019, Thermo Fisher Scientific), 67 ng total mRNA, 2.1 pmol total gRNA, and 0.38 pmol total annealed splint and donor DNA. After 16h, medium was changed back to DMEM with 10% FBS and cells were cultured until 2 days post transfection. For BFP to GFP conversion experiments, cells were dissociated with 0.25% trypsin, mixed with culture medium for trypsin inactivation, and analyzed with Attune NxT Flow Cytometer (Thermo Fisher Scientific). When multiple mRNAs, lmgRNAs, pegRNAs, or splints and donor DNAs were used in a single transfection, each was added in an equal amount to reach the total dosage described. When nicking gRNAs were used, they were added at half the dosage of the lmgRNA or pegRNA to reach the total dosage described.

### Primary cell transfection

PHH, PCH, and PMH were transfected 2 days after thaw with 0.3 uL Lipofectamine MessengerMAX (LMRNA015, Thermo Fisher Scientific) for co-delivery of all RNA and DNA components. Per well, the transfection mix contained 150 ng nCas9 mRNA, 150 ng ligase mRNA, 100 ng lmgRNA, 50 ng ngRNA, 5 ng splint, and 10 ng donor unless otherwise stated for optimization experiments. For pL-PGI transfections the amount of each splint and donor was halved to result in the same total dosage. PE control conditions contained 300 ng nCas9-RT fusion and 100 ng pegRNA, 50 ng ngRNA, or 100 ng each dual atgRNAs. Cells were taken down 3 days post transfection for analysis. For integration experiments, cells were co-transfected with 200 ng Bxb1 mRNA and either 1e6 multiplicity of infection (moi) AAV or 1000 moi HDAd 2-3 days post attB placement according to previously described procedures and taken down after an additional 5 days of culture.

### Genomic DNA extraction

For HEK293T cells, genomic DNA was extracted 2 days after transfection by removing medium, resuspending cells in 30 uL QuickExtract (QE0905T, LGC Biosearch Technologies), and incubating at 65C for 15 min followed by 98C for 10 min. For all primary cells, gDNA extraction was followed by magnetic bead cleanup (A63882 AMPure XP, Beckman Coulter).

### Target amplification and next-generation amplicon sequencing (NGS)

Target regions were amplified with Q5 Hot Start High-Fidelity Master Mix (M0494X, NEB) for 25 cycles using annealing temperatures for the gene-specific part of the primers calculated by NEB’s online tool (https://tmcalculator.neb.com/). Amplified targets were either imaged with gel electrophoresis in a 2% agarose gel or used as a template for the Illumina barcoding PCR 2.

### Sanger sequencing

Primers were designed to amplify a 500 – 800 bp region surrounding the edit site and target amplification was performed using Platinum Superfi II Master Mix (12368010, Invitrogen) using manufacturer recommended protocol. Samples were sequenced by Genewiz (Azenta) and analyzed using Synthego inference of CRISPR edits tool (https://ice.synthego.com/#/).

### In vitro toxicity

Toxicity of transfection conditions in cells was measured by CellTiter-Glo 2.0 (G9241, Promega) following manufacturer’s methods using 100 uL of premixed reagent per well in 96 well white-walled plates (165306, Thermo Fisher Scientific). Luminescence was detected on GloMax Discover Microplate Reader (GM3000, Promega).

### Droplet digital polymerase chain reaction (ddPCR)

Custom primers and probes were designed to measure editing in all referenced loci. Results were normalized to custom reference assays targeting unedited regions of the same genes in the respective species. Probes were dual labelled with 3′-3IABkFQ and either 5′-carboxyfluorescein (FAM) for edit targets or 5′-hexachloro-fluorescein phosphoramidite (HEX) for reference. Assays were validated using gBlocks representing edit outcomes to test for both specificity and linearity. All primers, probes, and gBlocks were synthesized by IDT. Each reaction contained 12 µL of 2x ddPCRSupermix for probes (No dUTP) (1863025 Bio-Rad), 1.2 µL of each primer and probe mix to final concentration of 0.5 uM for each primer and 0.25 uM for each probe, 0.12 µL each of HindIII and Eco91I (FD0505 and FD0394, Thermo Fisher Scientific), 10-20 ng of DNA and water to a final volume of 24 µL. Droplets were generated on the AutoDG Instrument for automated droplet generation (186410, Bio-Rad). PCR amplification was performed with the following cycling parameters: initial denaturation at 95 °C for 10 min, followed by 40 cycles of denaturation at 94 °C for 30 s and combined annealing/extension step at 58 °C for 1 min, and a final step at 98 °C for 10 min. Data acquisition and analysis were performed on the QX200 Droplet Reader.

### *In vivo* guide synthesis

Guide RNAs were synthesized on an AKTA Oligosynt synthesizer (Cytiva). Base-loaded 2000 Å CPG was packed into a 6.3 mL stainless steel column for synthesis scale of ∼50 umol. After solid phase synthesis, the oligonucleotide on support was treated with 15 mL of AMA and incubated for 4 hr at 25 °C. The solution was filtered, cooled on dry ice, and 15 mL of triethylamine trihydrofluoride was added dropwise. The reaction was heated for 4 h at 45 °C, then cooled on ice, quenched with ∼10 volumes of water, and neutralized. The crude product was purified on an AKTA Avant system (Cytiva) using a PLRP-S 300 Å, 15-20 um column with mobile phases 25 mM DBAA in H2O and 50% ACN at 60°C. Sodium salt exchange and desalting of the final product was done by TFF on a Sartoflow Smart system, using a 0.14 m2 regenerated cellulose membrane with 10 kDa molecular weight cutoff.

### Lipid nanoparticle (LNP) formulation

Lipid nanoparticles were formulated using the Precision Nanosystems Ignite. Nucleic acid payloads were diluted in a 50 mM pH 4.5 acetate buffer. All lipids were purchased from commercial vendors: ALC-0315 (Broadpharm), DSPC (NOF America), cholesterol (Avanti), and DMG-PEG2000 (NOF America). Lipids were diluted in ethanol and combined to create a final stock solution according to the desired molar ratio. Mixing was performed to achieve a final LNP composition with an N/P ratio=6. LNP were then diluted and dialyzed overnight. LNP were concentrated using ultracentrifugation and sterile filtered prior to dosing. All LNP were analyzed using DLS and Ribogreen to assess size, polydispersity, and encapsulation efficiency of cargo and determined to meet the desired QC parameters.

### *In vivo* mouse studies

All animal study procedures were approved by Explora BioLabs under IACUC protocol EB17-004-302. Female CD-1 mice (6 to 8 weeks) were ordered from Charles River Laboratories. LNPs were intravenously injected at a dose of 6 mg/kg calculated based on RNA content with 5 animals per treatment group. On day seven post LNP injection, animals were euthanized and liver tissue was collected from the median lobe from each animal for gDNA extraction (Quick-DNA/RNA Zymo Kit #R2131). ddPCR analysis performed following same methods as cell studies with efficiencies reported relative to mouse Tfrc reference copy assay (Applied Biosystems 4458366).

### Other

All figures were made with BioRender.com or Adobe Illustrator 2024. Graphs and data analysis were performed with GraphPad Prism 10. All results are reported as mean and standard deviation of biological duplicates or triplicates for *in vitro* experiments and individual results from 5 replicates with standard deviation for *in vivo* experiments.

---

**Extended Data Figure 1.**
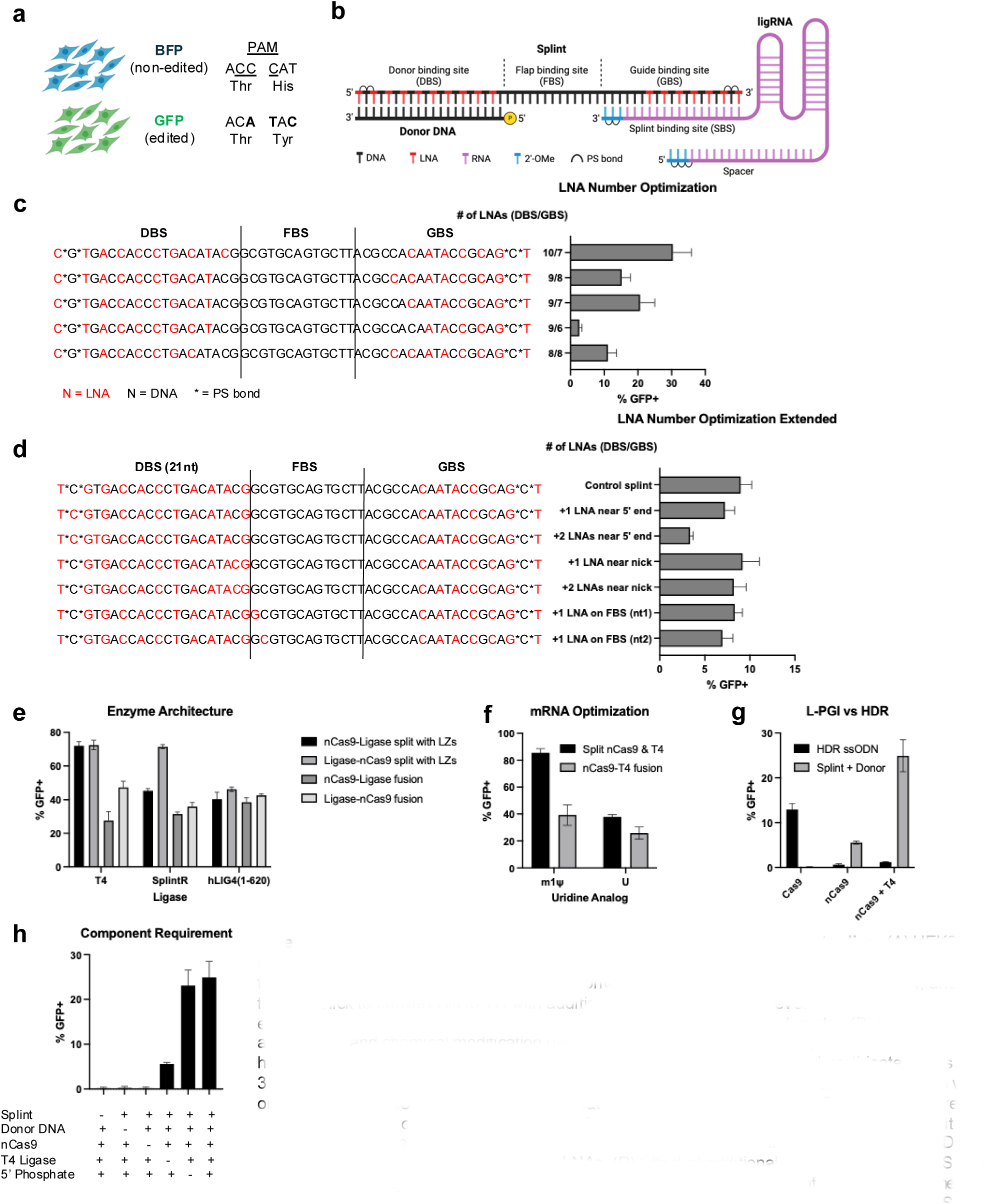
Additional optimization data in HEK293T reporter line. (A) HEK293T cells were virally transduced to express blue fluorescent protein (BFP) that would convert to green fluorescent protein (GFP) upon successful conversion of 3 nucleotides at positions +2, +4, and +5 from the nick to convert His to Tyr with additional PAM mutations to prevent repeat Cas9 engagement at the site. Conversion efficiency was assessed by flow cytometry. (B) Nucleic acid architecture and chemical modification pattern showing the lmgRNA, splint, and donor in their hybridized form. The donor contains a 5’ phosphate modification and phosphorothioate bonds at the 3’ terminus. The splint contains alternating DNA and LNA bases in the DBS and GBS regions with only DNA in the FBS and two phosphorothioate bonds at each end. The lmgRNA includes three 2’-OMe modifications on the 5’ and 3’ ends and phosphorothioate bonds at both termini. (C) Edit efficiencies using splints with the same base sequence but different numbers of LNAs in the DBS or GBS showing greater potency with more LNAs. (D) Effect of additional LNAs in the splint DBS and FBS. The control splint has alternating DNA and LNAs in the DBS. All splints include the same 19-nt GBS (not shown). (E) Additional enzyme format evaluation testing three ligases with nCas9. Split mRNAs all contained LZ placement at either the C or N terminus with corresponding placement on nCas9. Fusions tested either enzyme placement order with XTEN as linker. (F) Effect of pseudouridine (m1Ψ) in mRNA production in either a fusion or split mRNA system. (G) L-PGI mechanism verification using either a 100-nt ssODN donor for HDR or the L-PGI splint and donor combined with either Cas9, nCas9, or nCas9 + T4 ligase. (H) Dropout experiment demonstrating that all components are required except exogenous ligase supplementation and donor phosphorylation.

**Extended Data Figure 2.**
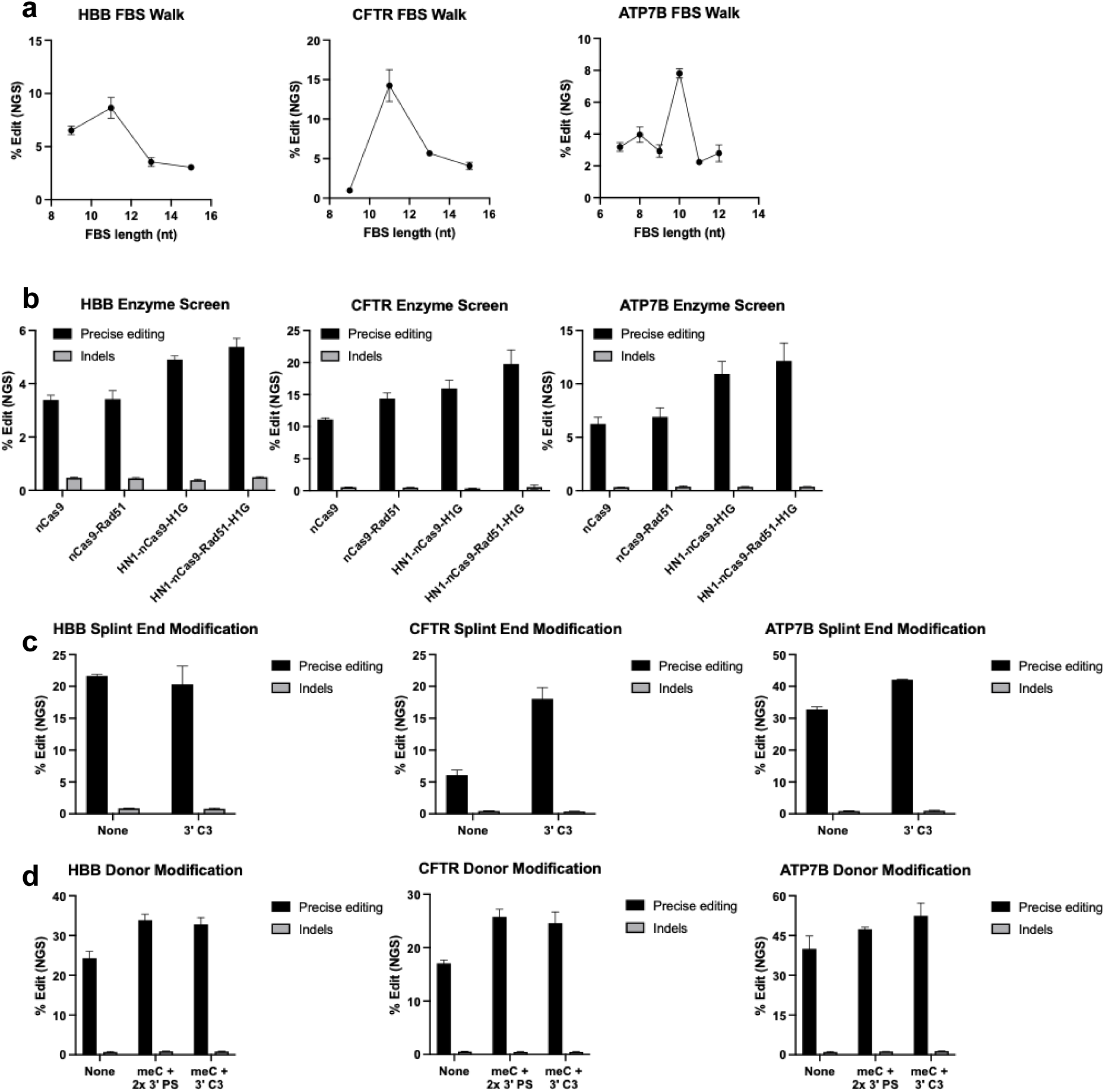
L-PGI optimizations for editing point corrections in 3 endogenous loci in HEK293T cells. All experiments used spacers identified in *HBB*, *CFTR*, and *ATP7B* for installing disease relevant mutations with added silent mutations to prevent sequential editing and 24nt donors with full splint-donor complementarity. Edits: *HBB* +4 A to T and +5 G to A (E6V, sickle cell disease), *CFTR* +4 T to C, +9 G to A and +14 C to T (R553X and G551D, cystic fibrosis), and *ATP7B* +5 G to T and +8 G to T (H1069Q, Wilson’s Disease). (A) Splint FBS length optimization in *HBB*, *CFTR*, and *ATP7B*. (B) Effect of DNA binding domains in nCas9 mRNA on edit efficiency and indel generation, all in combination with separate T4 ligase mRNA. (C) Editing efficiencies with or without a C3 spacer on the 3’ end of the splint for exonuclease protection. (D) Donor modifications showing impact of adding cytosine methylation (meC), phosphorothioate bonds (PS), and 3’ C3 spacer to improve affinity with the splint and protect against exonuclease activity.

**Extended Data Figure 3.**
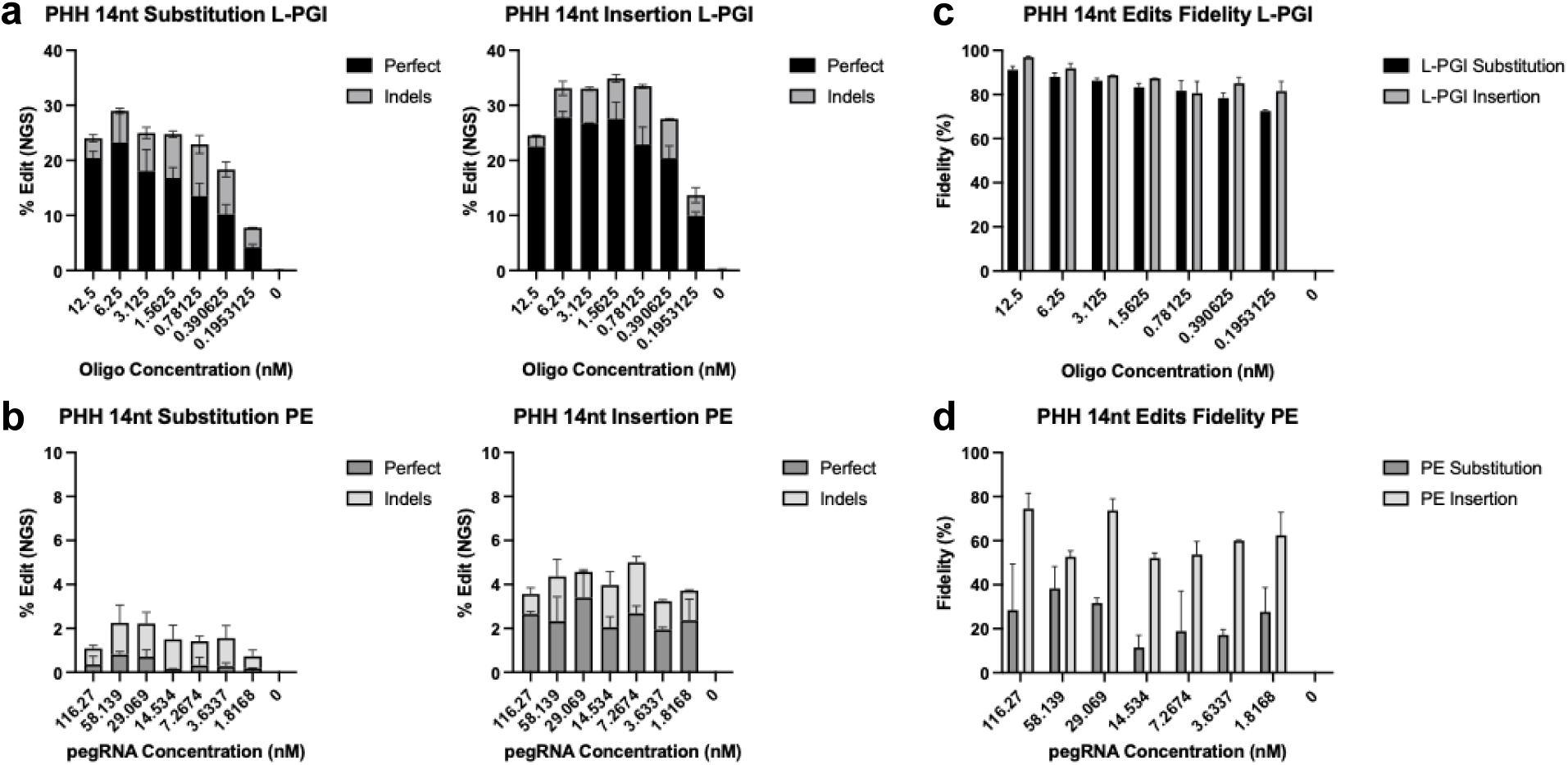
14 nt edit transfection optimizations with L-PGI and PE. (A) Perfect and indel efficiencies for 14 nt sequence substitution and insertion in *PAH* using L-PGI at titrated doses of splint and donor oligonucleotides showing concentration of each in cell culture media. All conditions used constant doses of other components formulated and co-delivered. (B) Perfect and indel edit efficiencies using PE consisting of fused nCas9-RT, pegRNA, and ngRNA using titrated doses of pegRNA with all other components held constant. (C) Effect of component ratio optimization on edit fidelity of L-PGI using varying splint and donor doses. (D) Effect of pegRNA dose optimization on fidelity for PE.

**Extended Data Figure 4.**
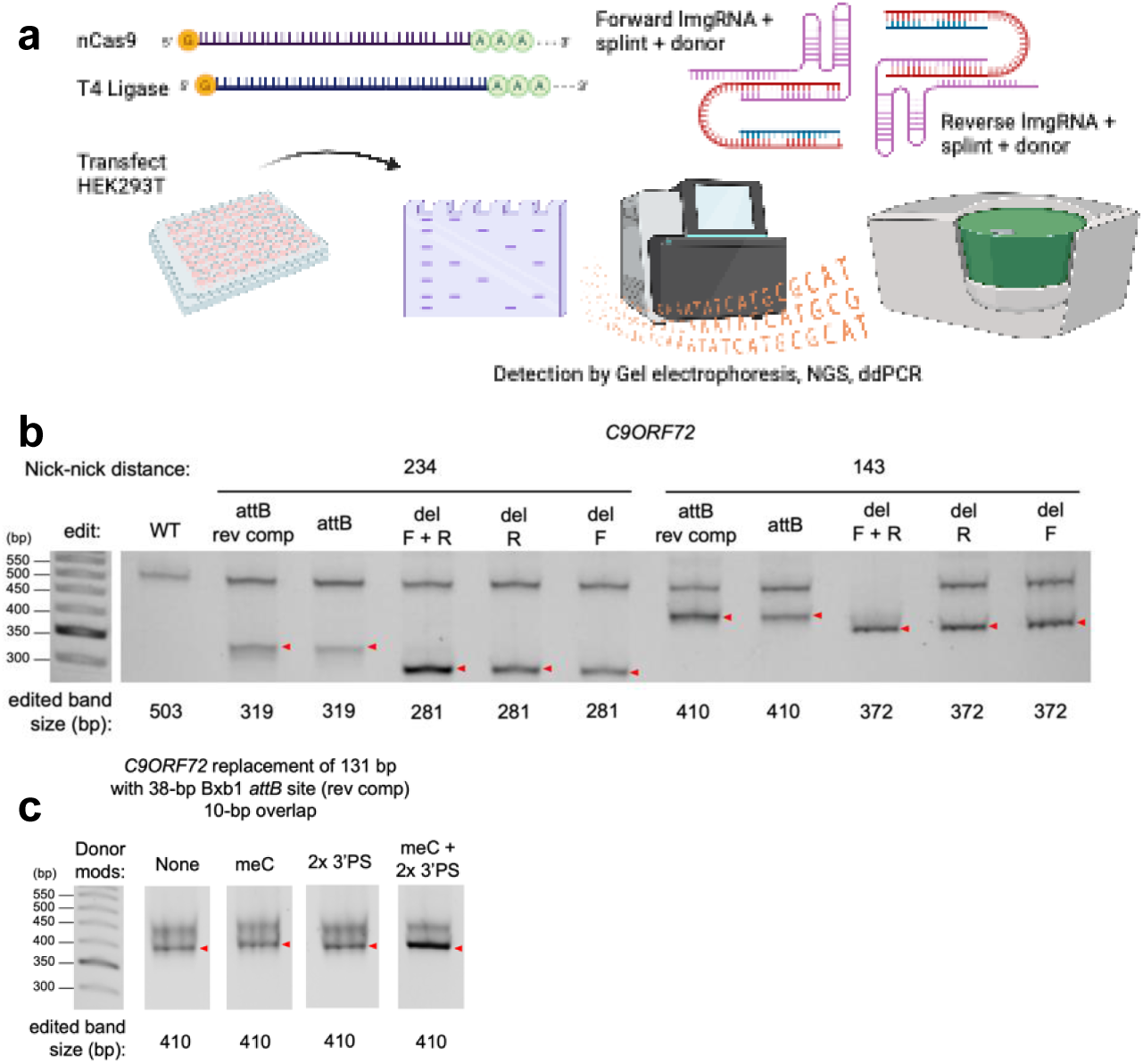
Schematic of pL-PGI transfection, detection methods and initial efficacy data. (A) Diagram of HEK293T workflow. Cells were co-transfected with IVT nCas9 and T4 mRNAs and chemically synthesized lmgRNAs, splints, and donors using Lipofectamine 2000 and MessengerMAX delivery. Cells were collected 72 hours after transfection, isolated for gDNA, and analyzed by either PCR and gel electrophoresis gel shift assay, NGS amplicon sequencing, or ddPCR with primers and probes for detection of insertion or excision at the target site normalized to wild-type reference allele copy number. (B) Agarose gel of PCR amplicons for *C9ORF72* after pL-PGI editing. The wild-type (WT) fragment is non-edited control and the fragments marked with red arrows indicate the desired replacement or deletion at the expected size. attB and attB rev comp are replacements of the nick-nick region with the Bxb1 attB site in forward or reverse orientation. del F + R is a deletion, del F is the deletion without the Rev splint and donor, and del R is the deletion without the Fwd splint and donor. (C) Agarose gel of PCR amplicons for the Bxb1 attB replacement edit of *C9ORF72,* comparing donor DNAs with different chemical modifications. The fragment of the desired edit is marked in red.

**Extended Data Figure 5.**
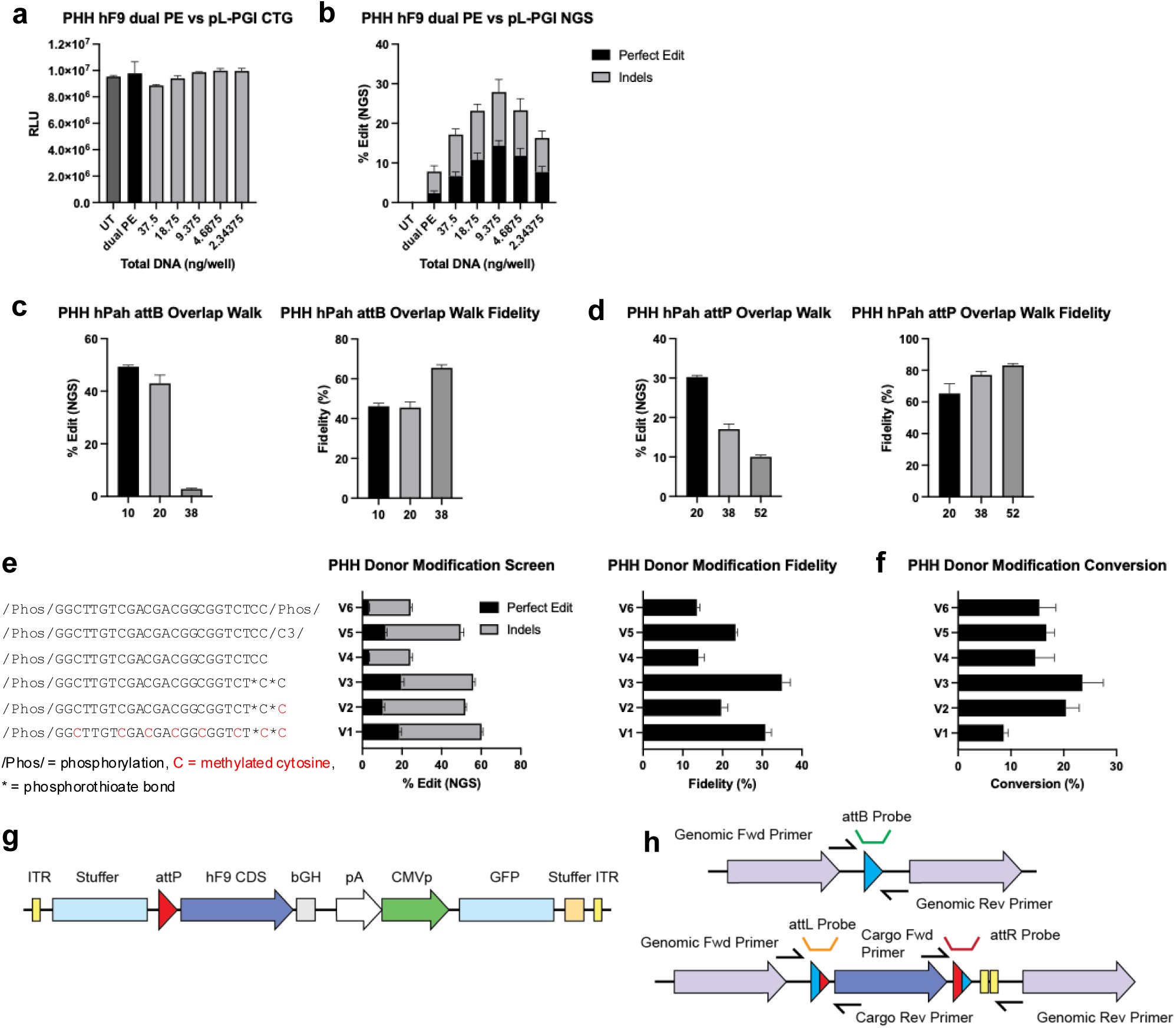
pL-PGI attB placement optimizations and Bxb1-mediated integration in PHH. (A) Assessment of tolerability at a range of splint and donor doses. PHH were transfected with either PE control consisting of nCas9 engineered RT fusion mRNA and dual attachment site guide RNAs (atgRNAs) or split nCas9 T4 LZ mRNAs and paired lmgRNAs with varying doses of equimolar splints and donors. Toxicity was determined by Cell Titer Glo (CTG) assay measuring relative luminescence (RLU) in cells 3 days post transfection. (B) Bxb1 38 bp attB placement in F9 intron 1 from the same experiment showing dose dependence of both efficiency and fidelity. (C) attB overlap walk in PAH symmetrically varying overlap length around the central dinucleotide to 10, 20, or 38 full overlap showing efficiency and fidelity. In contrast to F9 attB overlap walk (Figure 4a), in this experiment donor length varied independently of splint, resulting in 5 nt and 14 nt donor 3’ flap overhangs in 20 and 38 overlap respectively. (D) Similar overlap walk conducted for 52 bp Bxb1 attP with 20, 38, or 52 overlap showing efficiency and fidelity. (E) Impact of donor chemical modification on pL-PGI efficiency, correct editing, indel generation, and fidelity showing requirements for nuclease resistance. (F) attB conversion rates observed with the same donor chemical modification patterns. attB was placed in PHH using pL-PGI followed by self-complementary AAV LK03 attP cargo and Bxb1 mRNA co-transfection and collection for edit efficiency by ddPCR. (G) Schematic map of helper-dependent gutless adenovirus (AdV5) containing attP and F9 exon 2-8 coding sequence with GFP reporter and stuffer sequences totalling 30 kb linear cargo. (H) Schematic diagram of integration product and ddPCR primer and probe placement for specific detection of attB placement and integration over the attL or attR junctions. attB assay for detection of attB edited only control and unreacted residual attB: genomic forward primer, genomic reverse primer, and FAM labeled attB probe. attL assay: genomic forward primer, cargo reverse primer, and FAM attL probe. attR assay: cargo forward primer, genomic reverse primer, and FAM attR probe. All assays were run separately on the same biological samples and allelic edit efficiencies were calculated against a reference assay placed in nontarget PAH intron.

**Extended Data Figure 6.**
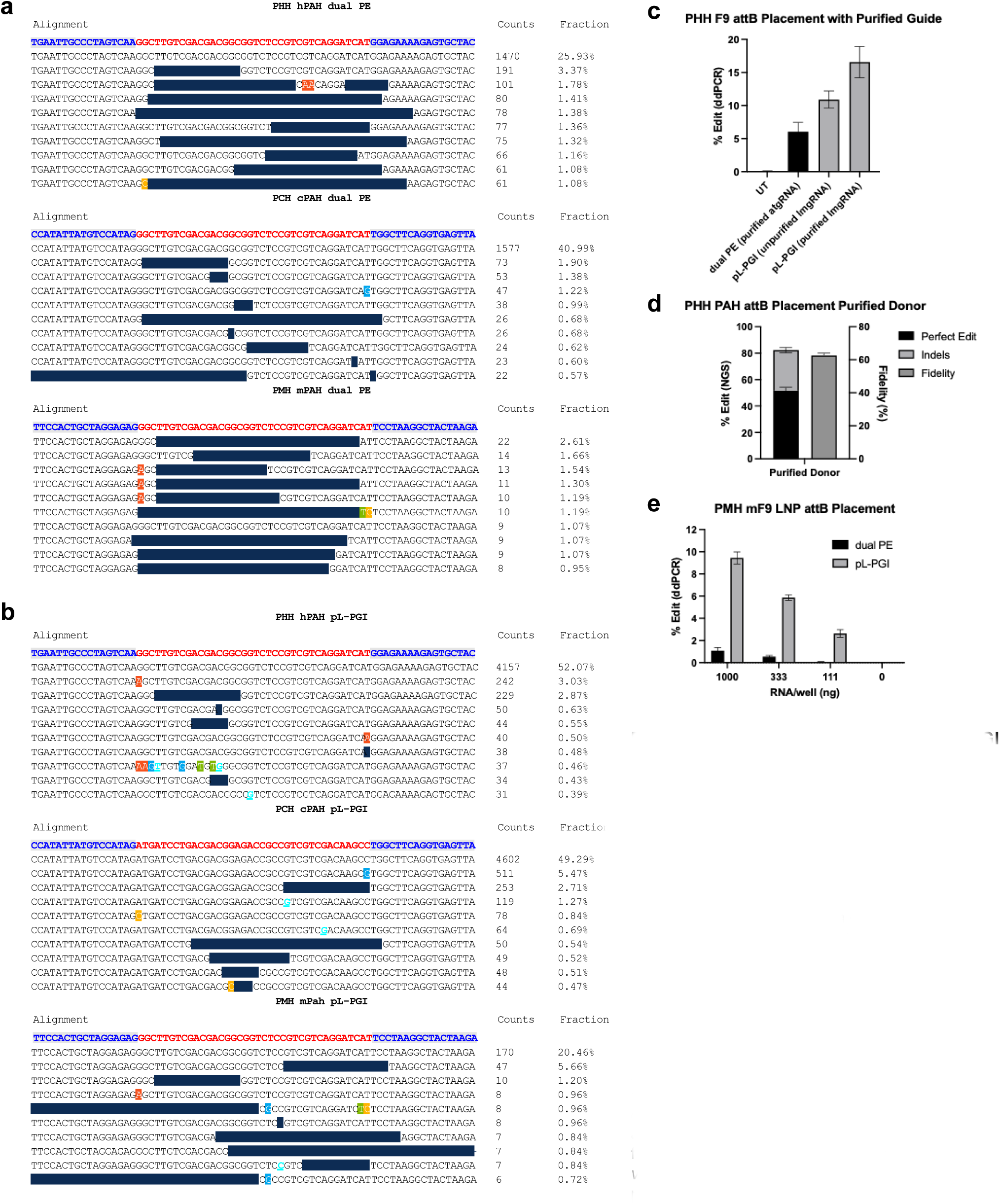
Comparison of pL-PGI with PE across species in vitro and additional data for in vivo translation. Sequencing alignments comparing PE (A) and pL-PGI (B) attB placement outcomes in PAH intron 1 in PHH, PCH, and PMH. Reference sequence is shown in blue and red differentiating genomic context and attB sequence (PCH pL-PGI contains reverse complementary attB). Sequencing results are presented below reference in each set showing only the top 10 reads sorted by abundance. In both PHH and PCH I the dominant read is perfect edit while PMH show dominance of excision and partial transcription. pL-PGI outcomes show greater quality and uniformity across species, with major imperfect products consisting of asymmetric attB insertion containing only forward or reverse 24 nt donor. (C) PHH attB placement efficiencies in *F9* with HPLC purified lmgRNAs or crude desalted lmgRNAs versus PE2 with HPLC purified atgRNAs. (D) PHH attB placement efficiency and fidelity in *PAH* with HPLC purified donor. (E) *In vitro* potency of AIO LNP delivering either PE or pL-PGI for attB placement in *F9* intron 1 of PMH.

